# Predicting CIN rates from single-cell whole genome sequencing data using an *in silico* model

**DOI:** 10.1101/2023.02.14.528596

**Authors:** Bjorn Bakker, Michael Schubert, Ana C.F. Bolhaqueiro, Geert J.P.L. Kops, Diana. C.J. Spierings, Floris Foijer

## Abstract

Chromosomal instability (CIN) drives the formation of karyotype aberrations in cancer cells and is a major contributor to intra-tumour heterogeneity, metastasis, and therapy resistance. Understanding how CIN contributes to tumour karyotype evolution requires quantification of CIN rates in primary tumours. Single-cell sequencing-based technologies enable the detection of karyotype heterogeneity, however deducing the actual CIN rates that underlie intra-tumour heterogeneity is still complicated. We have developed an *in-silico* model, called *CINsim*, to simulate the karyotype dynamics and validated our model in a murine mouse model for T-cell lymphoma (T-ALL) in which CIN is introduced by mutation of the Mps1 spindle assembly checkpoint protein. *CINsim* can simulate karyotype evolution within physiologically relevant timescales, across a range of CIN rates, and across a range of karyotype-imposed survival and proliferation effects. We find that *CINsim* can accurately predict the CIN rates in chromosomal instable mouse T-ALLs as well as in human colon cancer organoids as observed by live-cell time-lapse imaging. We conclude that *CINsim* is a powerful tool to estimate CIN rates from static single-cell DNA sequencing data by finding the most likely path from euploid founder cell to a heterogeneous tumour cell population.

## Introduction

Chromosomal instability (CIN) is a condition in which cells display an increased frequency of chromosome mis-segregation events in mitosis, and a hallmark feature of many cancers [1–4]. CIN will lead to cells with an abnormal DNA content, a state called aneuploidy [5,6]. The terms aneuploidy and CIN are often used interchangeably, but refer to different phenomena. Cells can be aneuploid without exhibiting CIN, resulting in a cell population of identical karyotypes [6,7]. Conversely, tumour cells that display a CIN phenotype will produce populations with cell-to-cell variability between karyotypes, termed karyotype heterogeneity. As CIN enables the rapid loss and gain of tumour suppressors and oncogenes respectively, it facilitates tumour cell evolution and is associated with metastasis, immune evasion, and chemotherapy resistance [1,2,8,9]. The prognostic value of CIN is further emphasized by the fact that copy number heterogeneity driven by CIN rather than (point) mutational heterogeneity correlates with poor survival in non-small cell lung cancer [10]. The frequency of chromosome missegregations, *i.e.* the CIN rate, is another important determinant of cancer cell fate as low CIN rates are insufficient to drive cancer while very high CIN-rates can be tumour suppressive in mouse models for CIN cancer [11,12]. Therefore, methods that can estimate CIN rates in cancer cells are essential to improve patient risk stratification.

Single-cell whole genome sequencing (scWGS) platforms have made it possible to measure complete karyotypes of individual cells at high resolution [13–16]. Recent scWGS efforts from various labs are revealing that cancers indeed frequently display intra-tumour karyotype heterogeneity, a strong indication of ongoing CIN [15,17–24]. Importantly, time-lapse imaging of primary tumour cultures with matched scWGS confirms that intratumour karyotype heterogeneity correlates well with observed CIN rates [15,19]. While time-lapse microscopy still remains the golden standard to determine the rate of CIN [6,15,17], live-cell imaging is not possible in primary human cancers. Therefore, estimating CIN rates from scWGS data could be a useful alternative. Recent work has attempted to do just that, although the predictive power of the mis-segregation model was still limited [25].

While modelling karyotype evolution has been done before to understand the effects of CIN on population growth rates and karyotype selection [25–30], none of the previously reported methods simulated evolution towards karyotype landscapes as observed in individual tumors and inferred the accompanying CIN rate. Furthermore, some of the models explored karyotype dynamics on timescales between several hundred and a few thousand cell divisions, yielding theoretical population sizes or time parameters that greatly exceed any physiological limit.

The overcome these limitations, we developed *CINsim*, an algorithm that models karyotype evolution from an initial diploid or tetraploid karyotype in an single cell towards the karyotype landscape of a cancer cell population as observed in primary cancer samples assessed by single cell DNA sequencing. By modulating the rates of CIN, cell death and cell proliferation *in silico*, *CINsim* finds the optimal values for these parameters that together yield a karyotype landscape closest to the landscape observed in the primary tumour considering a physiological number of cells and cell divisions. These optimal values represent the shortest path to this endpoint karyotype landscape and thus represent the most likely CIN, cell death and proliferation rates in the primary tumour. We used *CINsim* to model karyotype evolution for chromosomal instable acute T-cell lymphomas (T-ALL) derived from a mouse model in which the spindle assembly checkpoint protein Mps1 is conditionally mutated, and for a panel of human colorectal cancers. We find that *CINsim* correctly infers CIN rates as quantified experimentally by live-cell time-lapse imaging for both tumour types. We conclude that *CINsim* is a powerful tool to model karyotype evolution in chromosomal instable cancers and can be used to predict CIN rates from scWGS data.

## Results

### Performing forward stochastic simulations for chromosome mis-segregations in *CINsim*

Chromosomal instability leads to karyotype evolution and intratumour heterogeneity. As measuring karyotype dynamics in developing tumors is challenging, earlier studies have used *in silico* approaches to simulate karyotype evolution to better understand how karyotypes evolve in a CIN background [29,30]. While these modeling studies succeeded in simulating karyotype dynamics over time, they did not consider selective pressures for particular karyotypes as observed in cancer, nor did they incorporate physiological limitations such as tumor latency and maximum cell numbers. We therefore developed a model, *CINsim*, to examine karyotype dynamics within biologically relevant boundaries. *CINsim* simulates karyotype evolution across a range of parameters, most notably the CIN rate (defined by the chance of missegregation, *p_misseg_*, see below) and karyotype-dependent cell division and survival rates, within a physiological timescale. The model assumes that the combination of these parameters that yields a karyotype landscape that most resembles the landscape observed in the primary tumour corresponds to the actual path of karyotype evolution (Fig. 1a), an approach that to our knowledge has not been attempted before.

**Figure 1.**
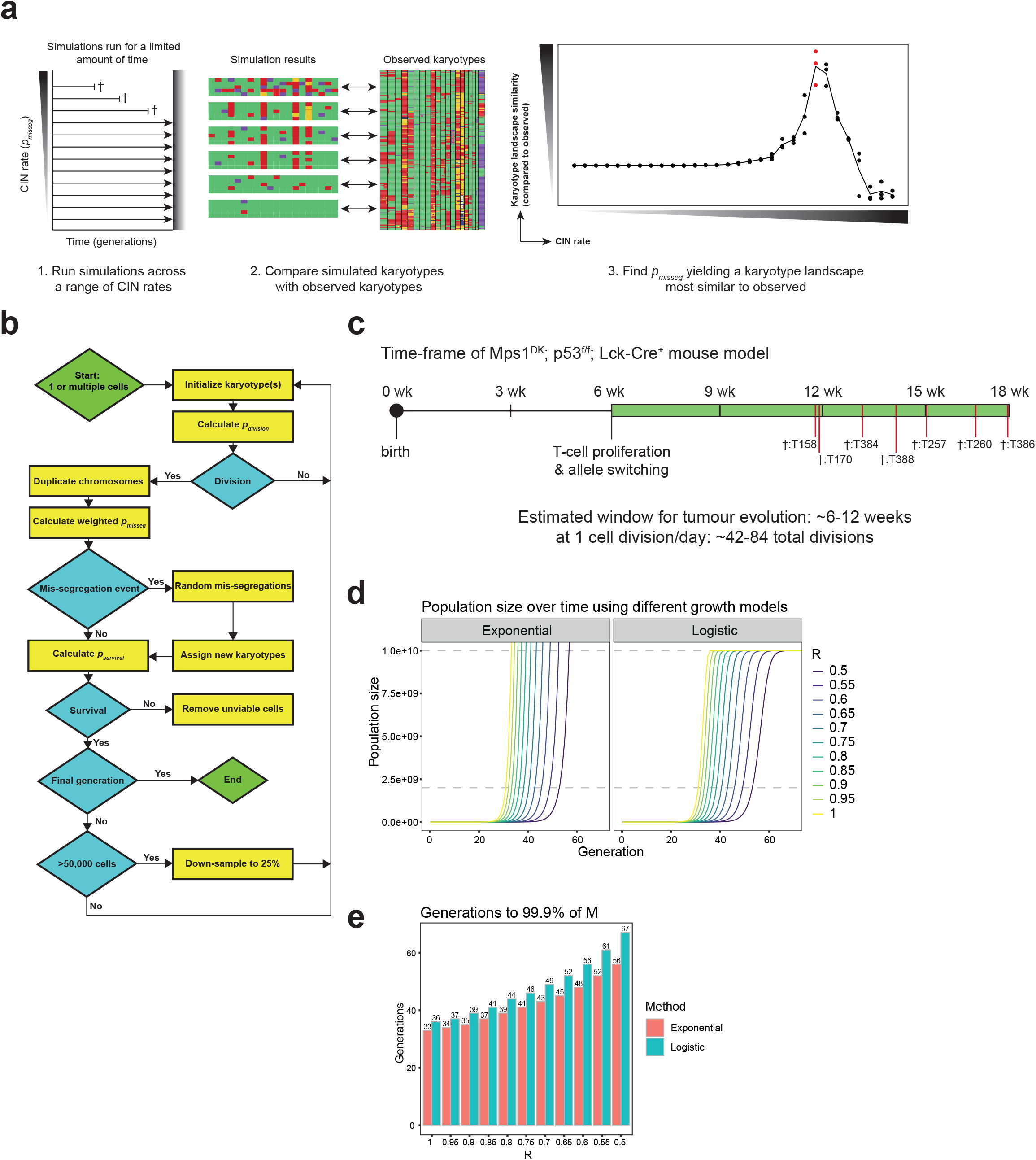
*CINsim* – a model for simulating karyotype evolution in an expanding cell population. **a.** Schematic overview of the workflow to identify the most likely CIN rate. Simulations are performed across a range of CIN rates, stopping at a predetermined set of generations or until the population reaches a certain size. Inferred karyotypes are then compared with observed ones, which is quantified using a similarity score. The value of *p_misseg_* that yields the greatest karyotype landscape similarity is selected as the most likely CIN rate. **b.** Schematic outline of the *CINsim* pipeline, with actions in yellow boxes and decision moments in blue diamonds. **c.** Schematic timeline for T-ALL development in the Mps1^DK^ mouse model. Red bars and crosses with tumour IDs represent the T-ALLs harvested at the indicated time points. The green bar represents the putative window of tumour evolution, estimated to be between 6 and 12 weeks. **d.** Expected population size over time at a range of division/survival rates *R* using either true exponential or logistic growth (with a carrying capacity at 10 billion cells). **e.** Required number of generations to reach 99.99% of carrying capacity *M* (10 billion cells) at a range of division/survival rates *R*. This provides a range of 30-70 generations in which to sample the karyotype landscape.

Within *CINsim*, cells are defined as single entities that each have a set of homologous chromosomes with their copy number represented by a single value, *i.e.* monosomy = 1, disomy = 2, trisomy = 3, etc.. Cell populations are represented in a table with chromosomes in columns and cells in rows. During one simulation cycle, cells divide into daughter cells and mis-segregate chromosomes according to a mis-segregation probability (*p_misseg_*) for each individual chromosome. Cells are removed after each simulated division if they contain nullosomies or more than 8 copies for a single chromosome, as these copy number states are exceedingly rare in scWGS data and hence likely not compatible with physiological constraints. Surviving cells undergo further selection by determining a karyotype fitness score for each individual cell). To determine a karyotype fitness score, chromosome copy number frequencies as observed in single cell whole genome sequencing (scWGS) experiments of primary tumour samples are used to determine the contribution to cellular fitness of each individual chromosome copy number state (*i.e.* frequently observed individual copy number states enhancing fitness and *vice versa*). Observed chromosome copy number state frequencies from scWGS data are used as scores, which are then summed across all chromosomes to yield a cell-specific karyotype fitness score. Karyotype fitness scores are then scaled into a probability of survival (*p_survival_*) according to the degree of selection (see Supplementary Methods). To determine whether a cell lives or dies, a random value between 0 and 1 is drawn for each cell from a uniform distribution. A cell ‘dies’ and is removed from the population when this value is larger than the *p_survival_* calculated for that cell. As such, cells that have a karyotype close to the most frequently observed karyotype in the populations and thus a *p_survival_* close to 1 will most likely survive, while cells with an infrequent karyotype will likely die. Simulation cycles are repeated until the estimated population exceeds a set threshold (discussed in more detail below), or until a pre-set number of generations is reached. To limit the required computational power, cell populations are downsampled to 25% whenever the simulated population size exceeds 50,000 cells. Figure 1b shows a schematic overview of the *CINsim* workflow and more details on the probabilities within the model, the effect of down-sampling on evolutionary dynamics, and estimating the true population size without down-sampling are described in the Supplementary Materials and Fig. S1. We quantified similarity between simulated and scWGS-observed karyotype landscapes using two methods, the karyotype measure similarity (KMS) and the Chromosome copy number Frequency Similarity (CnFS) score. This comparison is described in detail in the supplementary material under ‘quantifying the similarity between karyotype landscapes’ and revealed that the CnFS score performed best for our purposes (Fig. S2). The larger this CnFS value is, the better the simulation mimics the karyotype landscape observed in the primary tumour sample.

### Developing a model with physiologically relevant constraints

To test and optimise *CINsim*, we made use of prior-generated scWGS data from a mouse model for CIN-driven T-ALLs, in which the spindle assembly checkpoint protein Mps1 is conditionally mutated [17,31]. Based on this model, we first defined the physiological cell division and time constraints as observed for this mouse model. We limited the maximum number of cell divisions to 42-84 starting from a single cell towards full-blown T-ALLs as in this model T-ALLs take 6-12 weeks to grow from ~50 mg to ~1 gram with an average cell division time of 24 hours (Fig. 1c), [17,31]. In addition, we limited the maximum number of cells in an end-point tumour to 10 billion cells, assuming a single cell mass of 1 nanogram[32,33] and an end point tumour weight of 1-2 gram [17,31]. To get to 10 billion cells from a single cell, assuming exponential or logistic growth, requires 33 to 67 generations, respectively (Fig. 1d-e). Based on these physiological constraints, we limited the maximum number of *CINsim* simulation cycles to a 100 when modelling murine T-ALL, unless otherwise specified. These physiological restrictions form an important advance over prior models, which used much larger numbers (*i.e.* thousands) of generations for their simulations [26–30].

### Copy number-based selection yields convergence towards karyotypes observed *in vivo*

Chromosomal instability is a powerful driver of cancer cell evolution in which fitter cells are expected to thrive and unfit cells are selected against. As evolution assumes that the fittest individuals, or cells, will dominate the population [34], for *CINsim*, we assume that the frequency of a given chromosome copy number state observed in a tumour cell population is proportional to its effect on cellular fitness. Therefore, *CINsim* assumes that frequent karyotypes observed from scWGS data have a high chance of survival and infrequent ones do not. To model karyotype evolution in Mps1^DK^; p53^f/f^ Lck-Cre T-ALLs (from here onward referred to as Mps1^DK^ T-ALLs; DK refers to a truncation in the Mps1 kinase domain [31]) and to validate *CINsim*, we used a previously-generated scWGS dataset from 7 independent Mps1^DK^ T-ALLs (382 total single-cells; summary in Fig. 2a, all data in S3 [17]). In this model, recurrent karyotype features include trisomy 4/ 9, and trisomy or tetrasomy 14 /15 [31], while chromosomes 6, 7 and 8 are predominantly disomic. The remaining autosomes show varying degrees of aneuploidy, but are mostly disomic, in line with an ongoing CIN phenotype with little selection for a specific copy number alteration. Because our data includes both male and female tumours, aneuploidies for X are not considered.

**Figure 2.**
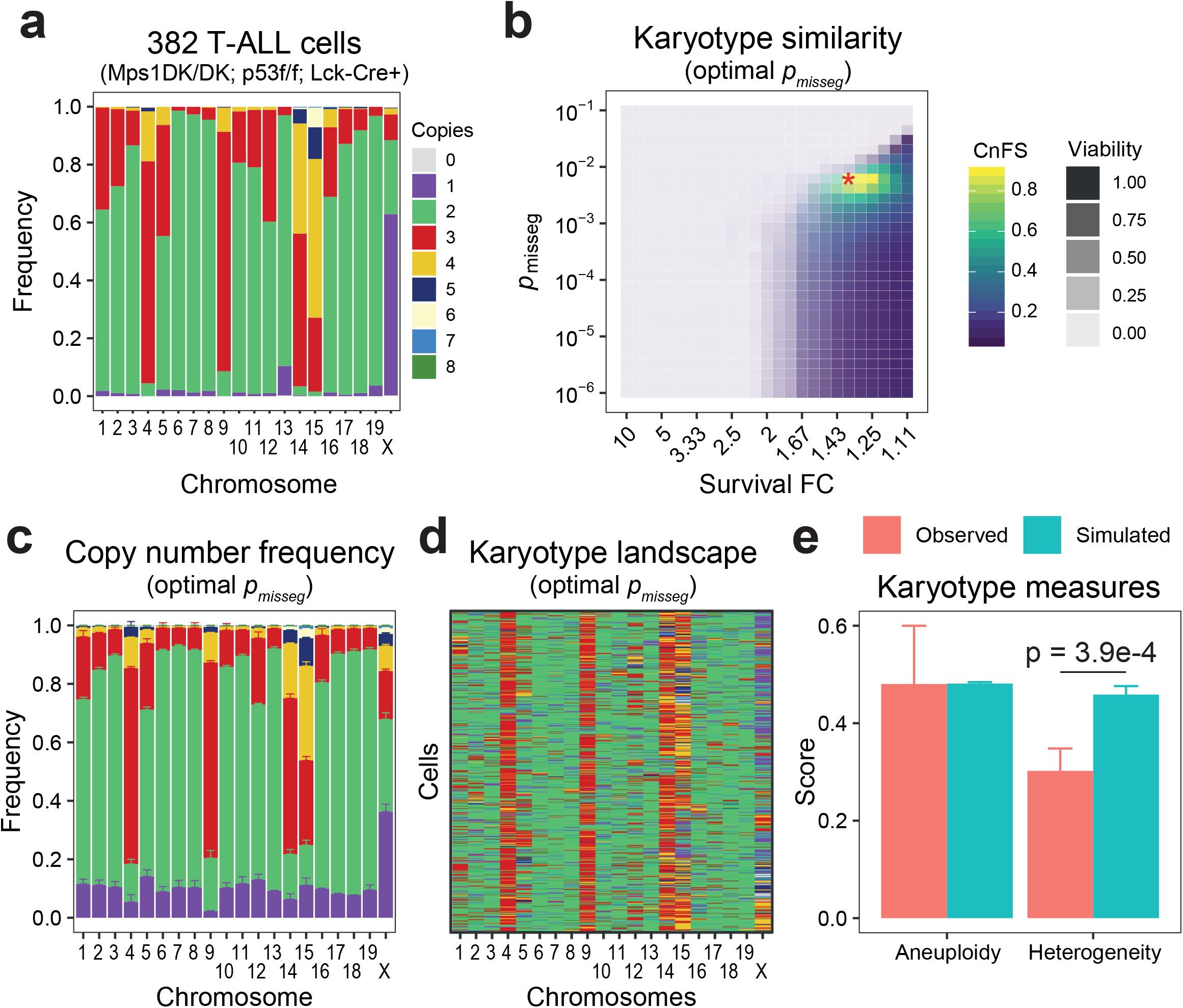
Chromosome copy number-based fitness results in rapid karyotype selection. **a.** Copy number frequencies in 382 T-ALL cells driven by Mps1 mutation and p53-loss as determined by scWGS of 7 independent lymphomas. **b.** Parameter scan for CIN rate (*p_misseg_*) and selection strength through survival (survival FC) for Mps1^DK^ T-ALLs using karyotype similarity as quantified by CnFS scores as the output. Red asterisk denotes the optimal combination of parameters yielding greatest similarity. Transparency corresponds to the proportion of simulations yielding a viable population (n = 100 iterations per parameter combination. **c.** Chromosome copy number frequencies at the *p_misseg_* yielding maximal CnFS. **d.** A representative example of 1,000 randomly sampled cells simulated by *CINsim* from the karyotype at optimal *p_misseg_* and Survival FC. **e.** Karyotype measures (genome-wide aneuploidy and heterogeneity scores) at optimal *p_misseg_* and Survival FC compared to observed measures. Non-viable simulations are included. Bars and whiskers indicate the mean±SEM. Significant differences in means was determined using a Wilcox signed rank test.

We then determined a ‘chromosome fitness score’ per individual chromosome, which is defined as the relative frequency of a given chromosome copy number state compared to the other copy number states for that chromosome in the scWGS dataset. For instance, if a chromosome would have a disomic state in 88% of the cells, a monosomic state in 11% of the cells and a trisomic state in 11% of the cells, the corresponding chromosome fitness scores would be 0.88 for the disomic state, 0.11 for the monosomic state and 0.11 for the trisomic state, respectively (Fig. S1e, top panel). The chromosome fitness scores were then combined into a ‘single cell karyotype score’, which represents the sum of all chromosome fitness scores for that cell. As such, cells with a high single cell karyotype score harbour karyotypes close to those observed in the scWGS dataset and thus represent ‘fit’ cells (Fig. S1e, bottom panel). Karyotype scores were then fitted to a probability of survival (*p_survival_*) such that aneuploid karyotypes yield a *p_survival_* ranging from 0.1 to 1.0, with euploid cells having a *p_survival_* of 0.9 assuming that aneuploid cancer cells can be more fit than euploid cells (Fig. S1f-g). The fold change in survival rate (survival FC) was then determined as the ratio of *p_survival_* of the daughter cell over the *p_survival_* of the mother cell. For instance, if the mother cell had a *p_survival_* of 0.9, and the daughter cell a *p_survival_* of 1 this yields a survival FC of 1.11, *i.e.* an increase of survival of 11%. When we generated 1,000,000 random karyotypes (with individual chromosome copy numbers restricted between 1 and 8), and near-diploid cancer-like karyotypes (with up to 4 aneusomies), we found that very few karyotypes yielded a greater karyotype score and *p_survival_* than euploid karyotypes (Fig. S1f), illustrating the narrow window for cancer cells to acquire karyotypes that are fitter than euploid cells.

Next, we simulated the proliferation of single founder cells in *CINsim* for a maximum of 100 generations at different CIN rates (*p_misseg_*) and rates of karyotype-dependent survival and compared the simulated karyotype landscape to the landscape as observed *in vivo* in Mps1^DK^ T-ALLs. At extreme CIN and selection rates (with *p_misseg_*>10^−2^ and survival FCs between 10 and 2) populations typically collapsed within 20 generations before karyotypes could evolved that would yield increased survival (Fig. 2b – light blue area). The reason is two-fold: firstly, populations with high CIN are either unable to stabilize their karyotypes on one that yields increased fitness, or evolve towards a lethal state. Secondly, the landscape of karyotypes yielding increased fitness is a subset of all possible and viable karyotypes, meaning cells are more likely to acquire a karyotype less fit than a euploid one. Enhanced selection will exacerbate the loss of fitness, equally resulting in rapid population extinction. However, at survival FCs of 2 and smaller we observed evolution beyond 20 generations, best resembling our scWGS data at survival FC 1.36 and *p_misseg_* 6.21×10^−3^. We found that for *p_misseg_* < 10^−3^ little to no adaptation takes place, whereas for *p_misseg_* ≥10^−2^ karyotypes will drift towards lethal nullisomies, leading to rapid population extinction (Fig. 2b). CIN rates between these values were compatible with evolution towards karyotypes resembling those of Mps1^DK^ T-ALLs, with a maximal similarity CnFS score of 0.897 (Fig. 2b, yellow area).

While the chromosome copy number frequencies simulated by *CINsim* were similar to those observed *in vivo* as measured by scWGS (compare Fig. 2a to Fig. 2c)., the simulated heterogeneity scores from *CINsim* were somewhat higher (Fig. 2e). In addition, the number of cells required to reach this level of karyotype landscape similarity after 100 generations were still well beyond physiological limits, with the average population size exceeding 10^10^ cells. We therefore conclude that while copy number-based selection enables the evolution of karyotypes similar to those observed *in vivo*, too many unfit cells are not yet selected against by *CINsim.*

### Karyotype-dependent survival and division rates both shape the karyotype landscape

To simulate the contribution of individual karyotypes to cancer cell proliferation, we converted karyotype scores into a division probability (*p_division_*) similar as done for *p_survival_*. As this alteration leads to a fraction of cells skipping a cell division each simulation cycle, we increased the maximum number of cycles to 250, or until 1010 cells were simulated, whichever requirement was met first. We then applied a range of *p_division_* fold changes (division FC), with the karyotype score only affecting *p_division_*. We found that as the division FC decreased, simulations required fewer simulation cycles to reach 10^10^ cells, and greatest similarity to our scWGS data was achieved at a division FC of 2.73 (Fig. 3a) at ~150 generations. Exchanging the *p_survival_* for a *p_division_* improved the CnFS score improved to 1.403 (Fig. 3c; compare to Fig 2b, CnFS score of 0.897 for *p_survival_*). The simulated karyotype landscapes were indeed more similar to *in vivo* karyotypes (compare Fig. 2a to 3b), suggesting that karyotype-dependent division has a greater role in karyotype selection.

**Figure 3.**
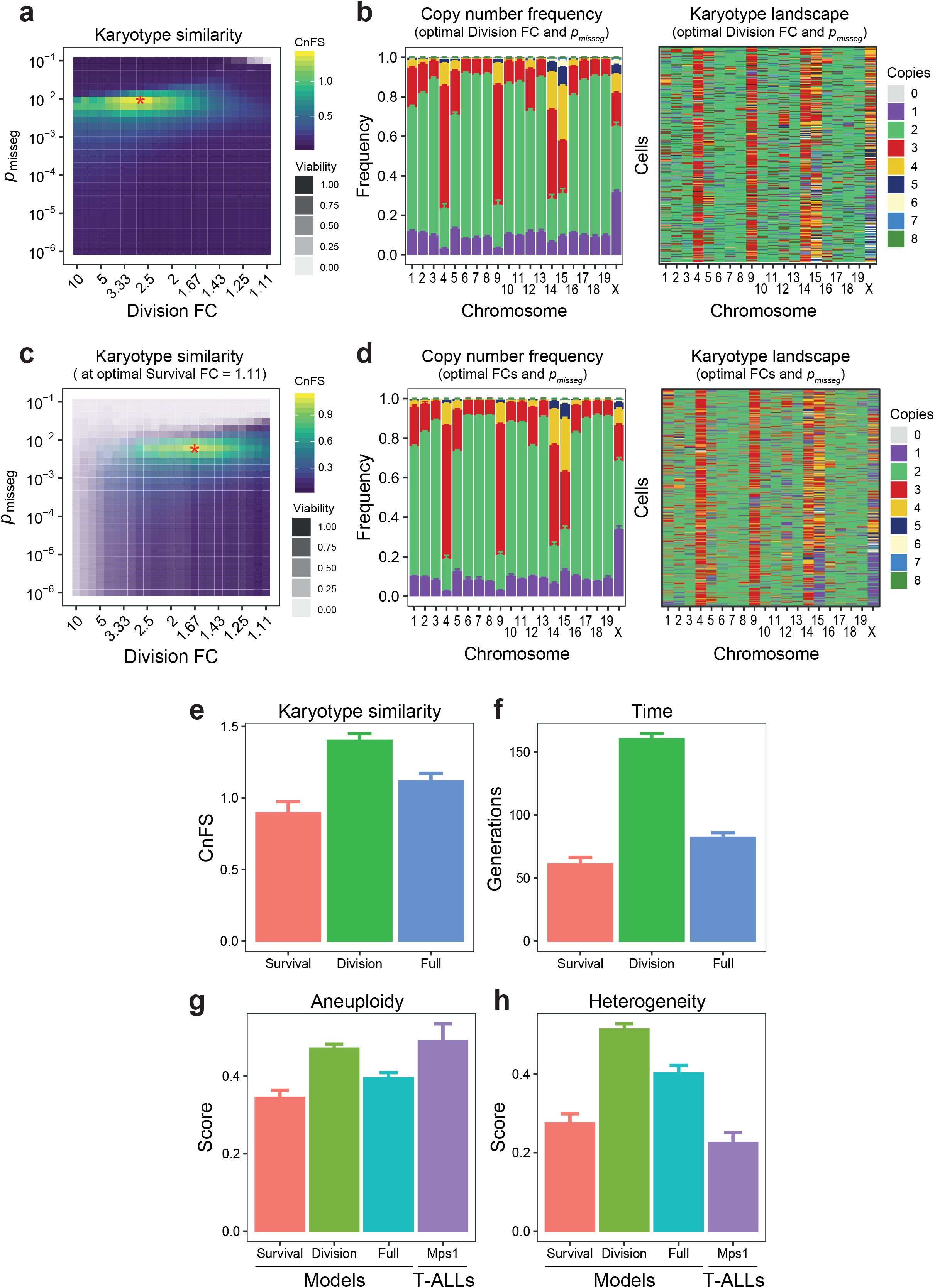
Karyotype-dependent survival and division rates shape the karyotype landscape. **a.** Parameter scan for CIN rate (*p_misseg_*) and selection strength through division (division FC) at optimal *p_misseg_* for Mps1^DK^ T-ALLs using karyotype similarity as output as quantified by CnFS scores. Red asterisk denotes the optimal combination of parameters. Transparency corresponds to the proportion of simulations yielding a viable population (n = 100 iterations per parameter combination). **b.** Chromosome copy number frequencies (left) and a representative example of 1,000 randomly sampled cells (right) at optimal parameters simulated by *CINsim* for the karyotype-dependent division model. **c.** Parameter scan for CIN rate (*p_misseg_*) and selection strength through division (division FC) at optimal *p_misseg_* and Survival FC for Mps1^DK^ T-ALLs using karyotype similarity quantified by CnFS scores as output. Red asterisk denotes the optimal combination of parameters. Transparency corresponds to the proportion of simulations yielding a viable population (n = 100 iterations per parameter combination). **d.** Chromosome copy number frequencies (left) and a representative example of 1,000 randomly sampled cells (right) at optimal parameters simulated by *CINsim* for the full model (karyotype-dependent division and survival). **e-h.** Maximal karyotype similarity (CnFS, **e**), time (generations, **f**) and karyotype measures (genome-wide aneuploidy, **g**; and heterogeneity scores, **h**) in viable populations compared between the three models and observed values in Mps1 T-ALLs.

We next explored whether combining both karyotype-dependent division and karyotype-dependent survival further improved the similarity between simulated and observed karyotypes, again restricting the number of simulation cycles to 250 and the population size to 10^10^. For this, we simulated karyotype evolution across a range of *p_misseg_* and division FCs at an optimal survival FC of 1.11, which was selected because it yielded the greatest CnFS scores on average (Fig. 3c). Combining optimized karyotype-dependent survival and division rates revealed that the optimal *p_misseg_* for the Mps1^DK^ model is around 6.21×10^−3^ corresponding to a mitotic error rate of 38.2% with a division and survival FC of 1.67 and 1.11 respectively (Fig. 3g). While combining all three parameters reduced the CnFS score to 1.119, overall, the simulated karyotype landscapes more closely resembled karyotypes *in vivo* (Fig. 3d), when also taking into account the number of cell divisions required and resulting aneuploidy and heterogeneity scores (Fig. 3e-g). Together, our simulations suggest that increasing the proliferation and survival rates of cells by 67% and 11%, respectively, at a chromosome missegregation rate of 38.2%, yields the most efficient route towards the karyotype landscape as observed in primary Mps1^DK^ T-ALL.

### Predicted CIN rates are concordant with rates observed in cultured murine T-ALLs

As *CINsim* predicted optimal CIN rates, we next compared these predicted CIN rates with the actual CIN rates observed *in vivo*. However, as quantifying chromosome missegregation rates in a developing thymic T-ALL is impossible with current available technology, we made use of a primary T-ALL cell line derived from the Mps1^DK^ model (T302; [17]). To rule out karyotype drift in culture conditions, we first single-cell sequenced our T302 cell line 3 passages after being taken into culture. Similar to primary T-ALLs, we found a preferential gain of chromosomes 4, 9, 14, and 15, suggesting that culturing conditions had minimal impact on the main karyotype distribution during the first few passages. In addition, chromosomes 1, 2, 5, 11, and 18 showed gains in the majority of cells (Fig. 4a & b), all of which are common copy number changes in our model (also see Fig. S3). Furthermore, aneuploidy and heterogeneity scores were very similar to those observed in primary T-ALLs, indicating that our primary cultures maintain their CIN phenotype during these early passages (Fig. 4c). We next used *CINsim* to simulate karyotype evolution towards the karyotype landscape we observed in cultured T302 T-ALL cells using scWGS similar across a range of division and survival FCs and CIN-rates (*p_misseg_*). We found that at division and survival FCs of 2.5 and 1.11, respectively, and a *p_misseg_* of 6.21×10^−3^ the simulated karyotype landscape most closely resembled that of primary T-ALL cultures (Fig. 4d). While clear selection occurred for chromosomes 1, 4, 9, 14 and 15, the simulated karyotype landscapes were more heterogeneous than observed in scWGS data (heterogeneity scores of 0.541-0.676 in simulated data vs. 0.276 in scWGS data; Fig. 4e & f). This discrepancy could be because the selection forces are underestimated, or the CIN rate is overestimated. To determine whether the *CINsim* predicted CIN rate was representative of the actual CIN-rates in this early passage T-ALL culture, we then quantified the chromosome mis-segregation rate in the T302 cell line using live-cell time-lapse imaging and found that 24% of mitoses show signs of unbalanced chromosome distribution (*i.e.* lagging chromosomes and anaphase bridges; Fig. 4g). This observed CIN rate is well in line with the *CINsim*-predicted CIN rate of 28.5-39.2% (Fig. 4g). We conclude that *CINsim* can simulate karyotype evolution as observed in our Mps1^DK^ T-ALL model and that *CINsim* can estimate chromosome missegregation rates reasonably well from copy number frequency data.

**Fig. 4.**
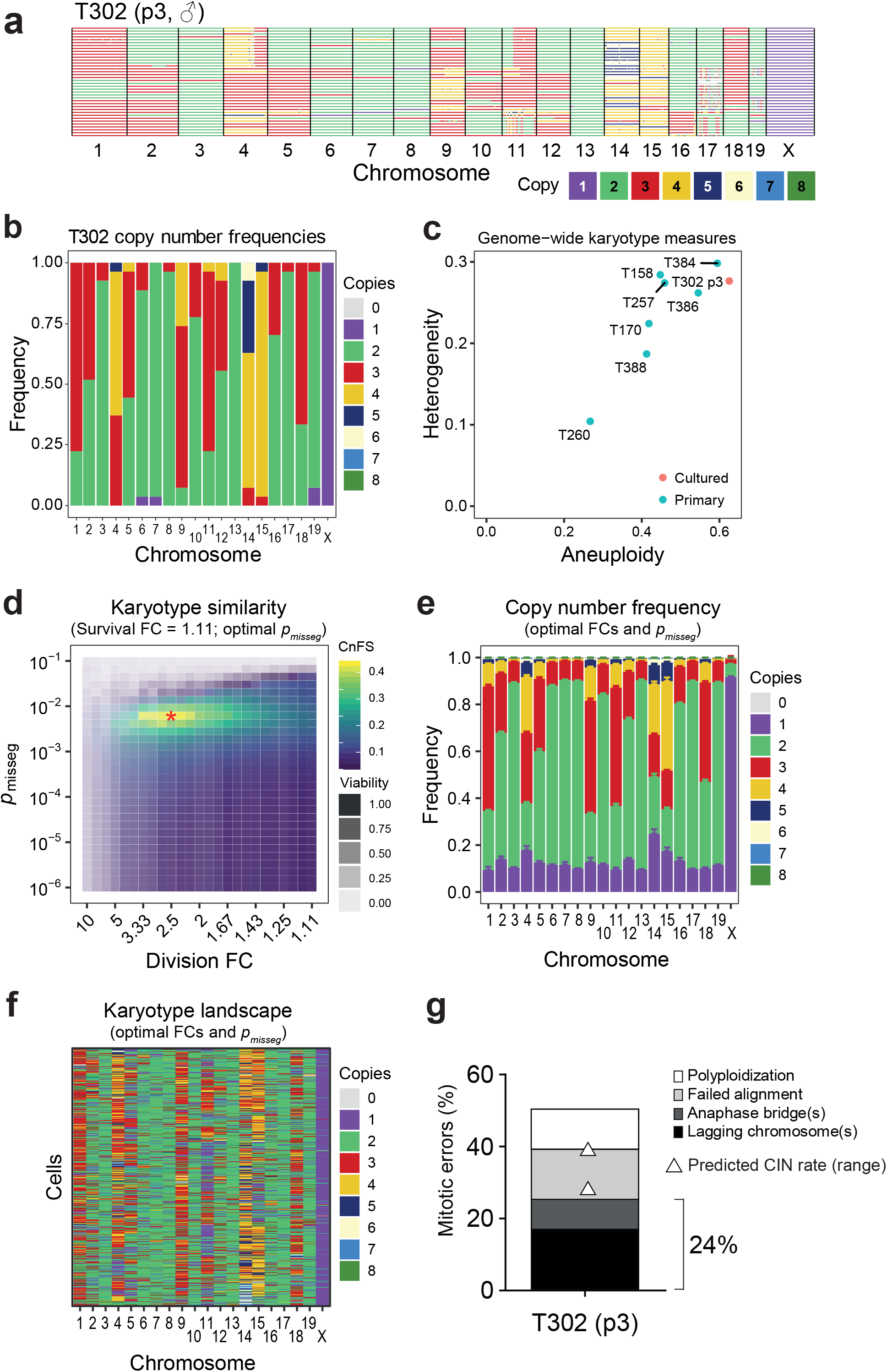
Predicted CIN rates validated by live-cell time-lapse imaging of cultured Mps1^DK^ T-ALLs. **a.** scWGS data from cultured T-ALL cells (cell line T302) after three passages. **b.** Copy number frequencies observed in T302 used for karyotype selection in *CINsim*. **c.** Karyotype measures (genome-wide aneuploidy and heterogeneity scores) for primary and cultured T-ALLs as calculated by AneuFinder. **d.** Parameter scan for CIN, Division and Survival FCs using karyotype landscape similarity quantified by CnFS as output for T302 simulations. Optimal parameter combination is indicated by a red asterisk. **e.** Copy number frequencies simulated by *CINsim* at optimal parameters. **f.** Representative copy number heatmap of 100 randomly sampled cells simulated by *CINsim* at optimal parameters. **g.** Observed mis-segregation rates in murine primary Mps1^DK^ T-ALL culture as determined by live-cell time-lapse imaging with predicted CIN rates in white triangles.

### *CINsim* predicts whole genome duplication as an early event in some, but not all CRC organoids

Finally, we wanted to validate *CINsim* for a completely different, human, cancer type. For this purpose, we re-examined scWGS data acquired from four human colorectal cancer (CRC) organoids displaying a range of aneuploidy and heterogeneity [15]. For this, we had *CINsim* model karyotype evolution based on the scWGS-observed karyotype landscape for each tumour organoid originating from a single diploid founder cells that proliferated for a maximum of 250 generations (assuming that human tumours take longer to grow than murine tumours) or until the population exceeded 5×10^10^ (the estimated carrying capacity for a ~4cm^3^ CRC tumour). Similar to our earlier simulations for murine T-ALL, *CINsim* yielded karyotype landscapes highly similar to those observed in the four tumour organoids (Figure 5a – left-side heatmaps, scWGS data in the centre column, similarity heatmaps in Fig. S4a, CnFS scores in Fig. 5b). However, for one of the tumour organoids (24TB, near-triploid karyotype), *CINsim* failed to predict the karyotype landscape.

**Figure 5:**
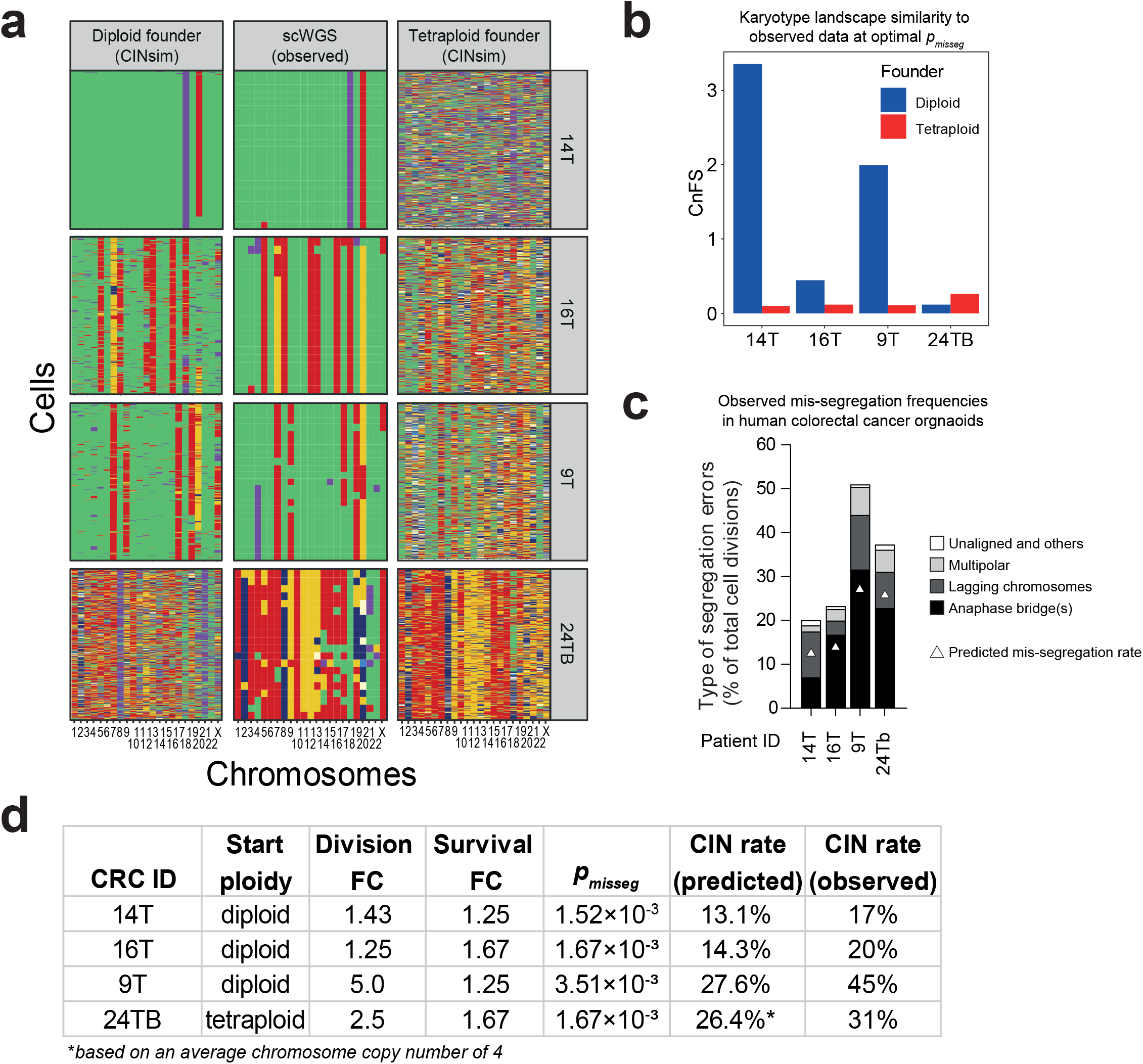
Predicting CIN rates in human colorectal cancers suggests that whole genome duplication is an early event during karyotype evolution. **a.** Karyotype landscapes at the optimal convergence of CIN and selection rates for four human CRC. *CINsim* simulations from diploid and tetraploid founders (left and right side, respectively) flank the observed karyotype landscapes as assessed by scWGS. **b.** Karyotype similarity as determined by CnFS scores for all four CRC organoids assuming diploid or tetraploid founders. **c.** Observed mis-segregation errors in human CRC organoids as determined by live-cell time-lapse imaging with predicted CIN rates in white triangles. Data adapted from [15]. **d.** Predicted founder cell ploidy, selection rates through cell division and survival, and mis-segregation frequencies as compared to observed CIN rates in organoid cultures.

Previous simulations studies have suggested that whole genome duplication (WGD) can help cancer cells to evolve while minimizing the chances for nullosomies, and that transitioning through the resulting tetraploid intermediate state favours evolution towards a near-triploid karyotype [30]. We therefore designed an additional simulation experiment for all four tumour organoids where we started from single tetraploid founders. Intriguingly, tetraploid founders allowed efficient evolution towards the optimal karyotypes for 24TB, but not for the near-diploid tumour organoids as reflected in the CnFS scores (Fig. 5a-b, similarity heatmaps in Fig. S4b). This suggests that of the four tumours only the near-triploid 24TB likely underwent whole genome duplication early during tumorigenesis.

Finally, we compared the optimal CIN rates as calculated by *CINsim* to those actually observed by live-cell time-lapse imaging of CRC organoids ([15], Fig. 5c-d). For tumour organoids 14T and 16T *CINsim* inferred a low CIN rate (13.1-14.3%) which is well in agreement with the observed values from time lapse imaging (17-20%), Similarly, organoids 9T and 24TB were predicted to have CIN rates of approximately 26.4-27.6%, which is reasonably close to the rates observed in culture (31-45%; Fig. 5c, d). Altogether, these analyses confirm that *CINsim* can be used to estimate actual CIN-rates from scWGS data for murine and human cancer and furthermore be used to determine whether cancer cells likely underwent whole genome duplication during tumorigenesis.

## Discussion

In this study we have developed a forward stochastic model, *CINsim,* to simulate karyotype dynamics using single cell karyotypes quantified by scWGS to infer CIN rates in primary tumours. Previous studies have explored arbitrary karyotypes yielding limited insights into the precise karyotype dynamics of tumours [25–30]. Since tumours are highly diverse in their degree of aneuploidy and karyotype heterogeneity, it is key to examine karyotype landscapes in individual tumours. Using our tumour-specific selection metric, we succeeded in simulating evolution towards a karyotype landscape that closely resembled the karyotypes observed in the primary tumours. We tested this approach on available data from several murine T-ALLs that were induced by mutation of the SAC protein Mps1 [17,31].

Unlike previously-published models, in *CINsim* simulations are restricted to a timescale and population size that are physiologically relevant for the sample of interest, thus giving cells a limited window in which they can evolve their karyotypes. By applying karyotype selection based on scWGS data from actual tumours, we succeeded in simulating karyotype landscapes highly similar to those observed *in vivo* and estimated optimal chromosome mis-segregation rates to achieve these landscapes. When accounting for both karyotype-driven effects on cell survival (*p_survival_*) and proliferation (*p_division_*), *CINsim* simulated karyotype landscapes with similar karyotype heterogeneity as observed by scWGS analysis in primary tumour samples and primary T-ALL cultures. Related to this, another recent modelling study using data from the same T-ALL models [35,36] that we used, also found that both cell survival and proliferation play an important role in shaping the karyotype landscapes of murine T-ALLs with a CIN phenotype (Ban et al., 2022).

The fact that *CINsim* could faithfully predict karyotype landscapes with similar aneuploidy and heterogeneity scores suggested to us that CIN rates in cultured T-ALL cells are in the same range as the CIN rates observed *in vivo*. Indeed, when we compared *CINsim*-inferred CIN rates to CIN rates observed by time lapse imaging, we found these to be in the same range for both T-ALL as well as CRC primary cultures, although observed CIN rates were ~2 fold higher inferred CIN rates for both primary cultures. A possible explanation is that observed mitotic normalities will only lead to karyotype changes in ~half of the cases as lagging chromosomes can also end up on the correct daughter cell. Therefore, these data strongly suggest that *CINsim* can infer CIN rates in primary tumours from scWGS data, an approach that, to the best of our knowledge, has not been attempted before.

Using live-cell microscopy remains the golden standard to quantify CIN rates [6]. To do so *in vivo* requires transgenic mouse lines with chromosome reporter constructs that allow for intravital time-lapse imaging. While such an approach is impossible in patients, culturing of primary tumours, tumour organoids or patient-derived xenografts is the next best alternative to yield insight into their CIN phenotypes. However, even this is still technically challenging, especially when studying (rare) tumour cells that are difficult or impossible to culture.

CIN is associated with tumour recurrence, increased chances of metastasis and therapy resistance, and thus an important determinant of patient prognosis [10]. However, different CIN rates might come with different prognoses. For instance, while high rates of CIN are found to suppress tumorigenesis, medium CIN rates are associated with more aggressive tumour development in mouse models [11,12,17], probably in a tissue specific manner. Therefore, faithfully assessing CIN rates in primary tumours might significantly improve treatment stratification. With rapidly decreasing costs of scWGS, *CINsim* could thus become an important diagnostic tool to estimate the CIN levels in tumour biopsies to help stratify treatment.

## Methods

### Basic characteristics of the *CINsim* chromosome mis-segregation model

For the first cycle in our simulations, we consider a cell to have a euploid karyotype (*i.e.* 40 chromosomes in a murine cell, 46 chromosomes in a human cell; with two copies of each chromosome, except for male sex chromosomes). Chromosomes have a defined copy number state (for euploid cells this number is 2) that will be inherited into two emerging daughter cells, unless a mis-segregation event occurs. The likelihood of a single chromosome copy mis-segregating is given by *p_misseg_* (hence *p*). Consider a chromosome pair, *i.e.* a copy number of 2 (diploidy). During S-phase both chromosomes are duplicated, leading to 4 sister chromatids. During mitosis, both duplicated chromosomes will be split and one copy for each duplicated chromosome will segregate into one of the daughter cells. Sister chromatids are bound unidirectionally by microtubules and pulled towards the nearest centrosome emitting the attached microtubule. A sister chromatid can be mis-segregated if it is instead pulled towards the opposite centrosome together with the sister chromatid (non-disjunction), or lags behind in the anaphase plane because of improper binding by microtubules (*e.g.* no binding or merotelic attachment). Given probability *p* of a single chromatid mis-segregating, the probability of no mis-segregation is 1 − *p*. More generally, the probability of any scenario of chromatids mis-segregating can obtained using the binomial theorem:

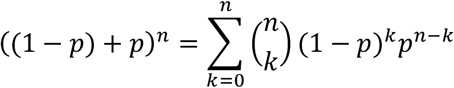

where *n* is the total of chromatids considered. For instance, for a chromosome with copy number 2, the probabilities of 0, 1 and 2 copies mis-segregating are given by (1 – *p*)^2^, 2*p*(1 – *p*) and *p*^2^ respectively:

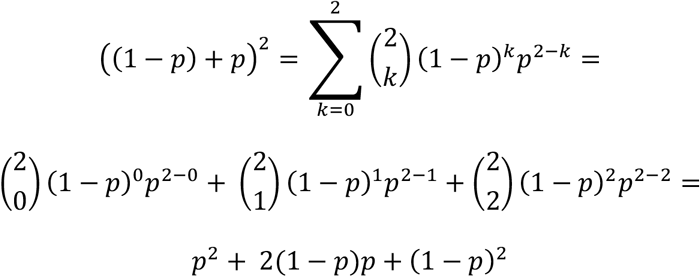

However, in this simplified example two simultaneous mis-segregation events (*p*^2^) do not result in disproportionate inheritance, as both mis-segregations cancel each outer out (*i.e.* both daughters acquire a single chromosome copy). In addition, as suggested by Laughney et al., *p*^2^ will be negligible for small values of *p*. From this, the probability of a mis-segregation event occurring is then given by 1 minus the probability of no mis-segregation: 1 – (1 – *p*)^2^. In this example the exponent of 2 is equal to the number of copies in the chromosome set at the time of mitosis onset. We can therefore generalize the weighted probability of any chromosome set mis-segregating (*p_weighted_*) to be:

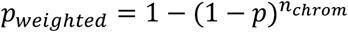

where *p* is *p_misseg_* and *n_chrom_* is the number of copies for that chromosome set at the time mitosis starts. This means that with increasing copy numbers a set has an increased probability of mis-segregating (Fig. S1A). To determine whether a chromosome set given *p_weighted_* mis-segregates in the simulation, a value is drawn from a uniform distribution (0, 1) and must be smaller than *p_weighted_*. Note that if *n_chrom_* is 0, *p_weighted_* will also be 0, as 1 − (1 – *p*)^0^ = 1 − 1 = 0, effectively forbidding the mis-segregation of non-existing chromosome sets. Also note that the mis-segregation probabilities of each chromosome set are independent of the copy number state of other chromosome sets. If a mis-segregation occurs, a single copy number gain and loss is then randomly assigned to the daughter cells. While in principle this method allows for the mis-segregation of multiple chromosome sets in one mitosis, single chromosome set mis-segregation events are more likely at physiological values for *p_misseg_*.

After every round of cell division, the viability of the individual daughters is assessed. In karyotype selection-neutral simulations, cells will die only if they lose all copies of a chromosome set (*n_chrom_* = 0) or exceed the maximum number of allowed copies (*n_chrom_* > 8). Viable cells then enter into a next round of cell division. In simulations with karyotype-based selection, these two boundaries still hold true, and the probability of cell survival is determined based on karyotype scores.

In the base model, all cells will undergo cell division simultaneously once per generation. To introduce asynchronous cell divisions, we have implemented a probability of division (*p_division_*) that can either be constant for all cells or be dependent on karyotypic fitness (see karyotype fitness below). Prior to cell division, a value is drawn from the uniform distribution [0,1] per cell. If the drawn value is greater than or equal to *p_division_* a cell will not enter mitosis but will enter subsequent selection based on copy number states and karyotype fitness. Individual cells are labelled at the start of the simulation with a unique identifier that progeny will inherit. This enables quantification of clonal abundance over subsequent generations when the founder population is greater than 1.

### Effects of down-sampling on simulation results and rates of evolution

Simulating millions of independent karyotypes requires a considerable amount of computational power and memory. The required time to simulate such large cell populations becomes impractical to study karyotype dynamics when adjusting many combinations of simulation parameters. As we also intend *CINsim* to be a tool that biologists can use on desktop computers or laptops to study simple evolutionary systems, we reduce the simulated population through random sampling whenever the total number of cells exceeds a particular threshold. However, the act of random down-sampling could result in the well-described ‘bottleneck effect’. Regular and substantial down-sampling of the population may therefore affect the rate of karyotype evolution. To test how strongly down-sampling affects simulation results, we performed simulations at various rates of down-sampling (a range of population size thresholds and down-sampling fractions). We allowed 200 non-mis-segregating clones to propagate synchronously for 50 generations (without the possibility of cell death) and determined the final fraction of clones still represented. We found that a down-sampling rate of 25% whenever the population exceeds 50,000 consistently results in 100% clonal survival while keeping simulation time per generation (Fig. S1B). These down-sampling parameters were therefore applied in all simulations.

### Estimating a biologically relevant timeframe for tumour evolution

Tumour cells in principle have the capacity to proliferate indefinitely, but are of course limited by availability of nutrients, oxygen, and space in their niche. In terms of population growth, tumour cells will therefore not display exponential growth but logistic growth instead. This environment-induced limit on the population size is termed the ‘carrying capacity’. In the described Mps1^DK^ T-ALL mouse model, this carrying capacity is the maximum number of cells that can exist in a single tumour, which we estimate to be 10^10^ cells. This is inferred from 1-2 gram tumours at an estimated lymphocyte weight of 1 ng [38], although this does not take into account possible dissemination from the original tumour site into peripheral blood or the bone marrow. Mathematically, the population size *P* at *t*+*1* given the size at *t*, and considering the carrying capacity of the environment, is defined as follows:

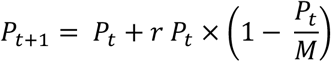

where *r* is the low-density growth rate (a balance between survival and death), and *M* is the carrying capacity. Starting from a single clone at *t* = 0 we calculated the number of generations required to reach 99.999% saturation at *M* for a range of values of *r* [0.5,1] reflecting different rates of tumour cell survival and cell division (Fig. 1d, e). We found that saturation is reached at a minimum of 30-34 generations and up to 52-69 generations for lower proliferation rates. This sets an estimated upper time limit for the development of murine T-ALLs between the order of 10 to 100 divisions.

### Simulation measures

After every round of cell division and selection we determine several population measures, including the survival rate, estimated true population size, and karyotype measures such as aneuploidy and heterogeneity scores based on AneuFinder [17]. The survival rate is defined as the fraction of daughter cells that survive after selection. Because of down-sampling the true number of cells is estimated from the simulated population size by multiplying with the down-sampling factor to the power of the down-sampling index:

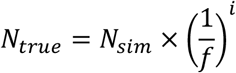

where *N_true_* is the true cell count, *N_sim_* is the number of simulated cells after one round of cell division and selection, *f* is the down-sampling factor (default of 0.25), and *i* is the down-sampling index (initial value is 0). The value of *i* will increase by 1 after each down-sampling event.

After selection, we calculate the degree of aneuploidy (*D*) and heterogeneity (*H*) of the population. These measures are based on the AneuFinder package[17]. For a population of *N* cells with *T* chromosomes, the aneuploidy score is defined as:

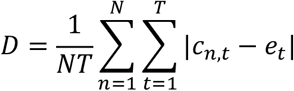

where *c_n,t_* is the copy number of chromosome *t* in cell *n*, and *et* is the euploid copy number of chromosome *t* (2 by default). The heterogeneity score is defined as:

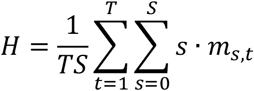

where *m_s,t_* is the number of cells with copy number state *s* for chromosome *t*, and *S* is the total number of copy number states present in the population. Importantly, *m_s,t_* is ordered for each chromosome, such that *m_s=0,t_* ≥ *m_s=1,t_* ≥ *m_s=2,t_*. This way a population with identical copy numbers for all chromosomes, whether they are considered aneuploid or not, will have a heterogeneity score of 0. In addition to the heterogeneity measure, we calculate the fraction of cells deviating from the modal copy number.

### Karyotype-based survival and division probability

To apply copy number-based karyotype selection we first made a matrix containing the observed frequencies of each chromosome copy number in 382 Mps1 T-ALL cells, with the copy numbers 0 to 9 in rows and the chromosomes 1 to X in columns. To ensure the X-chromosome does not affect karyotype fitness we assigned equal frequencies for the viable copy number states 1 to 8 (*i.e.* 1/8 = 0.125) and 0 to the lethal states (0 and >8). To determine the karyotype fitness score *S_f_* of a given karyotype, the chromosome copy numbers are matched to the corresponding chromosome copy number in the copy number frequency matrix obtained from scWGS data. For example, if chromosome 1 is observed at copy numbers 2 and 3 in scWGS at frequencies of 0.2 and 0.8 respectively, a karyotype within the simulation that has chromosome 1 at copy number 2 will score 0.2 for that chromosome, or 0.8 if chromosome 1 were at copy number 3. This process is repeated for all chromosomes, and all scores are summed into a single karyotype fitness score *S_f_*. In this way, cells with commonly observed copy numbers for certain chromosomes will obtain a greater fitness scores than cells with uncommon chromosome copy numbers. To assign a probability of survival to *S_f_* we first determined *S_f_* for 1,000,000 randomly generated near-diploid karyotypes (at most 4 to 5 aneuploid chromosomes) to ensure the scores follow a symmetric distribution (Fig. S1F). We next fitted these scores to the formula *a* × *Sf* + *b*, where *a* and *b* are the slope and intercept coefficients determined using the lm() function in R, such that euploid karyotypes had a probability of 0.9 (unless otherwise indicated in the text), and the highest possible score yielded a probability of 1. For this fit we found *a* = 0.04049, *b* = 0.4041 (Fig. S1G).

In summary, the survival or division probability of any karyotype is calculated as follows:

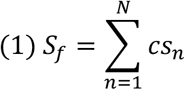

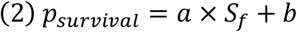

where *S_f_* is the karyotype fitness score, *N* is the total number of chromosome sets, *csn* is the assigned copy number score of chromosome *n*, and where *a* and *b* are coefficients with the fitted values as described above.

### Quantifying similarity between karyotype landscapes

When comparing multiple CIN rates and to determine at which rate the simulated karyotype landscape most resembles the observed landscape, we needed a metric that determines the similarity between the simulated landscapes. To this end, we developed two different metrics.

The first measure compares observed and simulated copy number frequencies. We define a score named copy number frequency similarity (CnFS) as the inverse sum of squares of differences between observed and simulated copy number frequencies:

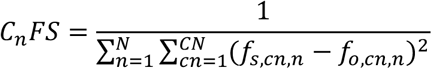

where *f_s,cn,t_* is the frequency for copy number state *cn* of chromosome *n* in simulated data, and *f_o,cn,t_* is the observed frequency for the respective copy number state of chromosome *n*. More dissimilar copy number frequencies will yield a greater denominator, resulting in smaller CnFS scores, and vice versa. Since the X-chromosome is selection-neutral (*i.e.* its copy number does not affect survival probability) in our simulations, we do not consider it to calculate karyotype similarity.

The second method is based on chromosomal aneuploidy and heterogeneity scores as they are determined by AneuFinder. We define a score named karyotype measure similarity (KMS) as the inverse sum of squares of differences between observed and simulated karyotype measures:

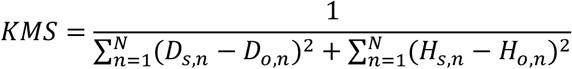

where *D_s,n_* and *H_s,n_* are the aneuploidy and heterogeneity score respectively for chromosome *n* in simulated data, and *D_a,n_* and *H_a,n_* are the observed karyotype measures. More dissimilar karyotypes will yield a greater denominator, resulting in smaller KMS scores, and vice versa. As with the CnFS metric, we do not include the X-chromosome when calculating the similarity score.

To test whether either one or both metrics adequately quantify differences between karyotype landscapes, we applied both metrics to three sets of karyotypes: 1) random near-diploid karyotypes, 2) karyotypes with preferential copy number gains for chromosomes 4,9, 14 and 15, and 3) karyotypes of Mps1^DK^ T-ALLs as determined by scWGS (Fig. S2A). Using both metrics through pairwise comparison we could cluster the karyotypes of the three sets together based on their similarity, with the random karyotypes clustering away from karyotypes that had preferential copy number changes (Fig. S2B).

We next tested how the two metrics change as karyotype evolution occurs, and whether it can be used to identify a karyotype landscape that best resembles the one observed in Mps1^DK^ T-ALLs. We ran simulations at a single rate of CIN (*p_misseg_* = 0.0025) for 250 generations, applying either no selection or Mps1^DK^-based copy number selection (see section on karyotype-based selection above), and determined the KMS and CnFS scores with the overall Mps1 T-ALL karyotype landscape as a reference (Fig. S2C). In simulations without selection both scores increased for the first 50 generations and then declined to baseline (CnFS) or even below baseline (KMS). In simulations with selection CnFS continued to go up as generations passed, whereas KMS score plateaued after 125 generations and afterwards decreased in some simulations (Fig. S2C). Surprisingly, the simulation with the highest KMS score had the lowest CnFS score, and vice versa. Monosomy 15, rather than trisomy/tetrasomy 15, was highly abundant in the simulation with the highest KMS score. In contrast, the top CnFS simulation showed karyotypes carrying trisomy and tetrasomy 15 (Fig. S2D). We found that the KMS metric did not consider the directionality of various aneusomies (*i.e.* monosmy and trisomy equally affect the aneuploidy score). Using the CnFS score we identified a landscape with the trisomy and tetrasomy 15, and additional aberrations commonly observed in Mps1^DK^ T-ALLs. For all simulations we therefore used the CnFS score to quantify karyotype landscape convergence.

### T-ALL culture, scWGS and time-lapse imaging

To acquire T-ALL cultures, Mps1^DK^ mice suffering from T-ALL were sacrificed when showing signs of lymphoma (weight loss, laboured breathing, and other signs of anaemia). Enlarged thymuses were dissected and homogenized through a 70 μm filter (Greiner) to acquire single cell lymphocyte suspensions. T-ALL cells were cultured in RPMI-1640 GlutaMax medium containing 25 mM HEPES (Gibco), 10% FBS (Sigma), 1% penicillin/streptomycin (Gibco), 1% non-essential amino acids (Gibco) and 55 μM β mercaptoethanol according to an established protocol [39]. Cells were passaged 1:10 when population density reached 2-3 million cells/ml, typically every 3 days.

For scWGS, T-ALL cells were harvested, sorted as single cells in 96 wells plates using a Jazz FACS flow cytometer (BD) into nuclear lysis buffer, processed in a semi-automated fashion to acquire single cell sequencing libraries and sequenced in a multiplex manner on an HiSeq sequencer (Illumina) as described previously [40]. scWGS data was analysed using AneuFinder [17].

For time-lapse imaging, primary T-ALL cultures were transduced with retroviruses carrying H2B-GFP [31] using spinfection. Transduced T-ALL cells were transferred onto a Lab-Tek imaging chamber (Nunc) and monitored using a 40x objective on a DeltaVision time-lapse microscope (Applied Precision/GE Healthcare/Leica). Time-lapse data were analysed using SoftWorx (Applied Preciesion) and ImageJ software and quantified in Excel (Microsoft).

## Code availability

All simulations were performed in R v3.5.0. The code for running *CINsim* and for making the figures is available from the GitHub repository at “bbakker1989/CINsim2”. A more detailed description of *CINsim* itself can be found in the methods section above.

## Acknowledgements

We thank the members of the Foijer lab for fruitful discussion. This work was supported by Dutch Cancer Society (KWF-kankerbestrijding) grants 2012-RUG-5549, 2015-RUG-11457 and NWO TOP grant 91215003 awarded to FF.

## Author contributions

B.B. and F.F conceived the project with assistance of M.S. B.B. wrote the code and performed all simulation and experiments. M.S. and R.W. co-supervised coding. D.C.J.S. supervised scWGS. A.C.F.B. and G.J.P.L.K. contributed data. F.F. supervised the project and provided funding.

---

**Table S1:** Characteristics of the murine Mps1^DK^ T-ALL panel.

| ID | Sample description | Sex | Age (wks) | Thymus weight (mg) | Spleen weight (mg) | Total tumour mass (mg) | # scWGS libs | Aneu ploidy binwise | Heterogeneity binwise | Aneu ploidy whole chrom | Heterogeneity whole chrom |
| --- | --- | --- | --- | --- | --- | --- | --- | --- | --- | --- | --- |
| T158 | Endpoint T-ALL<br>Mps1f/f; p53f/f; Lck-Cre+ | F | 11.4 | 1420.0 | 210.0 | 1630.0 | 88 | 0.4477 | 0.2841 | 0.449 | 0.269 |
| T170 | Endpoint T-ALL<br>Mps1f/f; p53f/f; Lck-Cre+ | M | 11.7 | 1280.0 | 0.0 | 1280.0 | 39 | 0.4832 | 0.2242 | 0.464 | 0.201 |
| T257 | Endpoint T-ALL<br>Mps1f/f; p53f/f; Lck-Cre+ | F | 15.1 | 1410.0 | 210.0 | 1620.0 | 86 | 0.4589 | 0.2742 | 0.453 | 0.249 |
| T260 | Endpoint T-ALL<br>Mps1f/f; p53f/f; Lck-Cre+ | M | 17.1 | 400.0 | 740.0 | 1140.0 | 46 | 0.3319 | 0.1042 | 0.323 | 0.102 |
| T386 | Endpoint T-ALL<br>Mps1f/f; p53f/+; Lck-Cre+ | F | 18.3 | 1780.0 | 50.0 | 1830.0 | 38 | 0.6048 | 0.2620 | 0.624 | 0.274 |
| T384 | 13 week T-ALL<br>Mps1f/f; p53f/f; Lck-Cre+ | M | 13.0 | 570.0 | 50.0 | 620.0 | 45 | 0.6579 | 0.2983 | 0.668 | 0.296 |
| T388 | 14 week T-ALL<br>Mps1f/f; p53f/+; Lck-Cre+ | M | 14.0 | 650.0 | 50.0 | 700.0 | 41 | 0.4770 | 0.1870 | 0.459 | 0.187 |
|  |  | Mean | 14.37 | 1072.9 | 187.1 | 1260.0 |  | 0.4945 | 0.2334 | 0.4914 | 0.2254 |
|  |  | SD | 2.63 | 526.2 | 257.6 | 470.7 |  | 0.1074 | 0.0686 | 0.1170 | 0.0672 |
|  |  | Total |  |  |  |  | 383 |  |  |  |  |
| T302 P3 | Cultured T-ALL (pas 3)<br>Mps1f/f; p53f/f; Lck-Cre+ | M | 16.7 | 590.0 | 60.0 | 650.0 | 38 | 0.6253 | 0.2765 | 0.675 | 0.268 |

**Figure S1.**
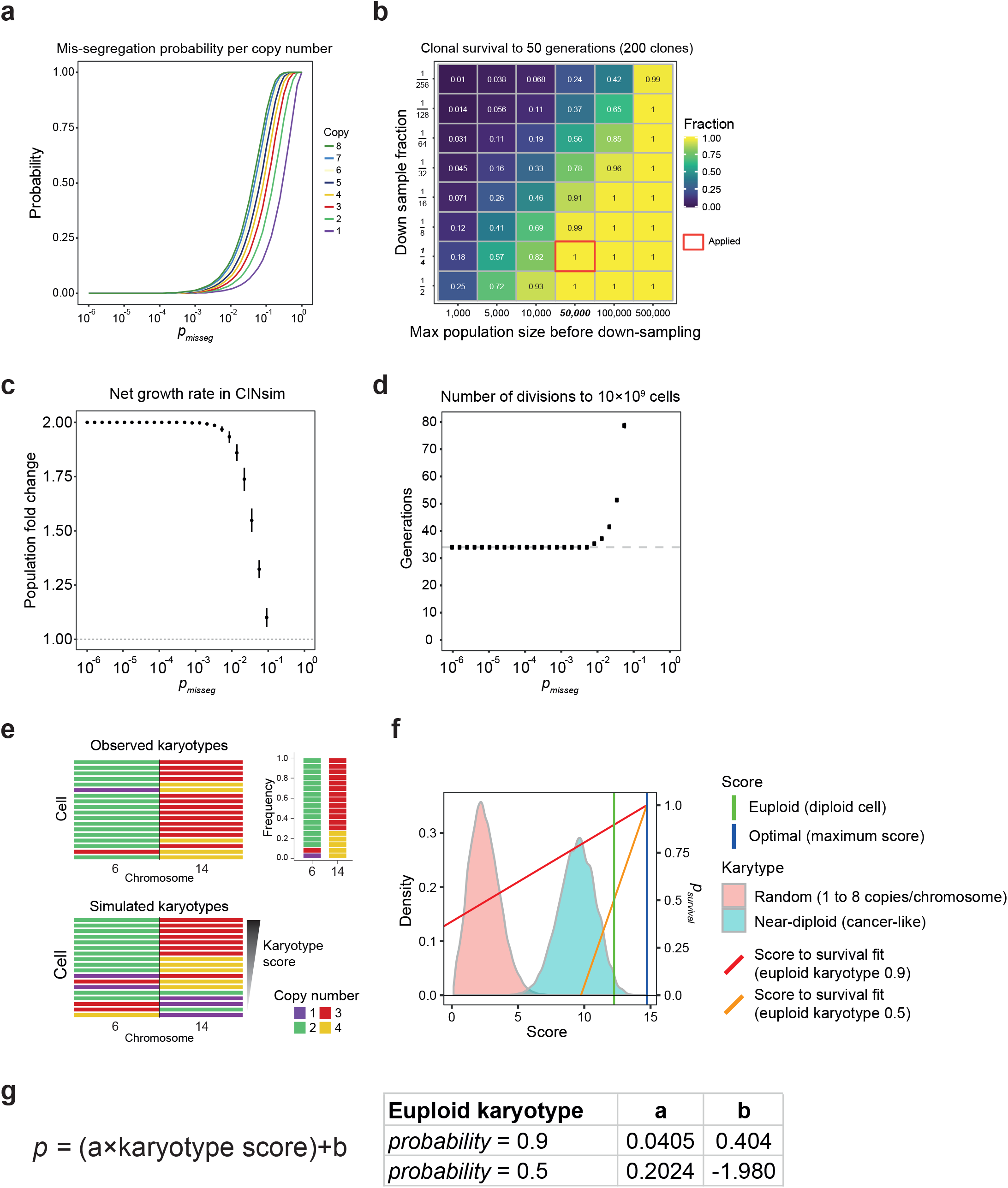
Basic properties of the *CINsim* model and karyotype-dependent selection. **a.** Expected mis-segregation probability per chromosome copy number over a range of CIN rates for a mouse genome. **b.** Optimisation of down-sampling strategy. 200 non-mis-segregating clones were expanded simultaneously, and down-sampling was performed over a range of population sizes and down-sampling fractions. At 25% down-sampling at a population size of 50,000 clonal survival is complete. These down-sampling parameters were applied in all simulations. **c.** Net growth rate, defined as the population fold change as a function of CIN rate (*p_misseg_*), demonstrating that high CIN rates are inherently lethal within the model. **d.** Time (in generations) required to reach 10 billion cells at different values for *p_misseg_*. Data is the same as in c, with the dashed line showing the theoretical minimum of 34 generations. **e.** Observed frequencies of copy number states for a chromosome are assumed to represent the most fit state for that chromosome. Cells with infrequently-observed chromosome copy numbers have lower karyotype scores than cells with frequently-observed copy numbers. **f.** Karyotype scores for a population of cells with either randomly generated (random copy numbers between 1 and 8 for any chromosome) or near-diploid (cancer-like) karyotypes, with the maximum possible karyotype score and euploid scores indicated in blue and green vertical lines respectively, highlighting that the majority of viable karyotypes are less fit than a euploid karyotype. The red line indicates the fit to convert a karyotype score into a survival/division probability such that a euploid karyotype corresponds to a survival probability of 0.9 (red) or 0.5 (orange). **g.** The formula used to convert a karyotype score into a probability, with coefficients shown for a euploid karyotype survival at 0.9 and 0.5 probability.

**Figure S2.**
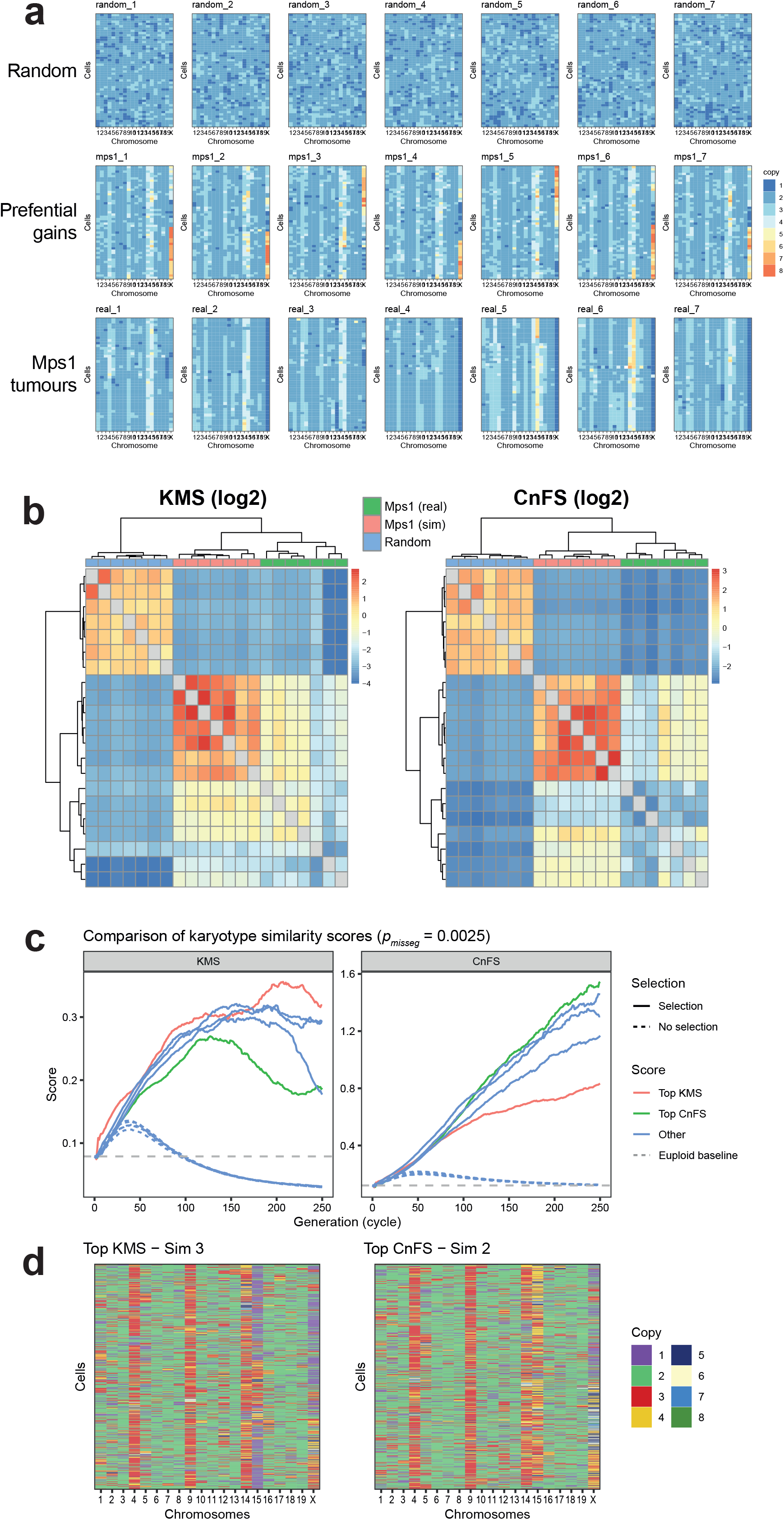
Comparing karyotype landscape similarity metrics, based on aneuploidy and heterogeneity (KMS) or chromosome copy number frequencies (CnFS). **a.** Heatmaps showing the karyotypes on which the two metrics are tested, either random near-diploid, or near-diploid with preferential copy number gains (+4, +9, +14, +15), and observed Mps1^DK^ T-ALL karyotypes. **b.** Karyotypes from panel a are compared paired-wise using the KMS and CnFS metrics, showing karyotypes from the same group clustering together, as do karyotypes with preferential karyotypes (simulated or Mps1^DK^ T-ALLs). **c.** Karyotype similarity to Mps1^DK^ T-ALLs for simulations at *p_misseg_* = 0.0025 over 250 generations. Top simulation according to either KMS or CnFS are indicated. Grey line indicates the baseline metric at the start of the simulation. **d.** Representative copy number heatmaps of the top simulations from panel c.

**Figure S3.**
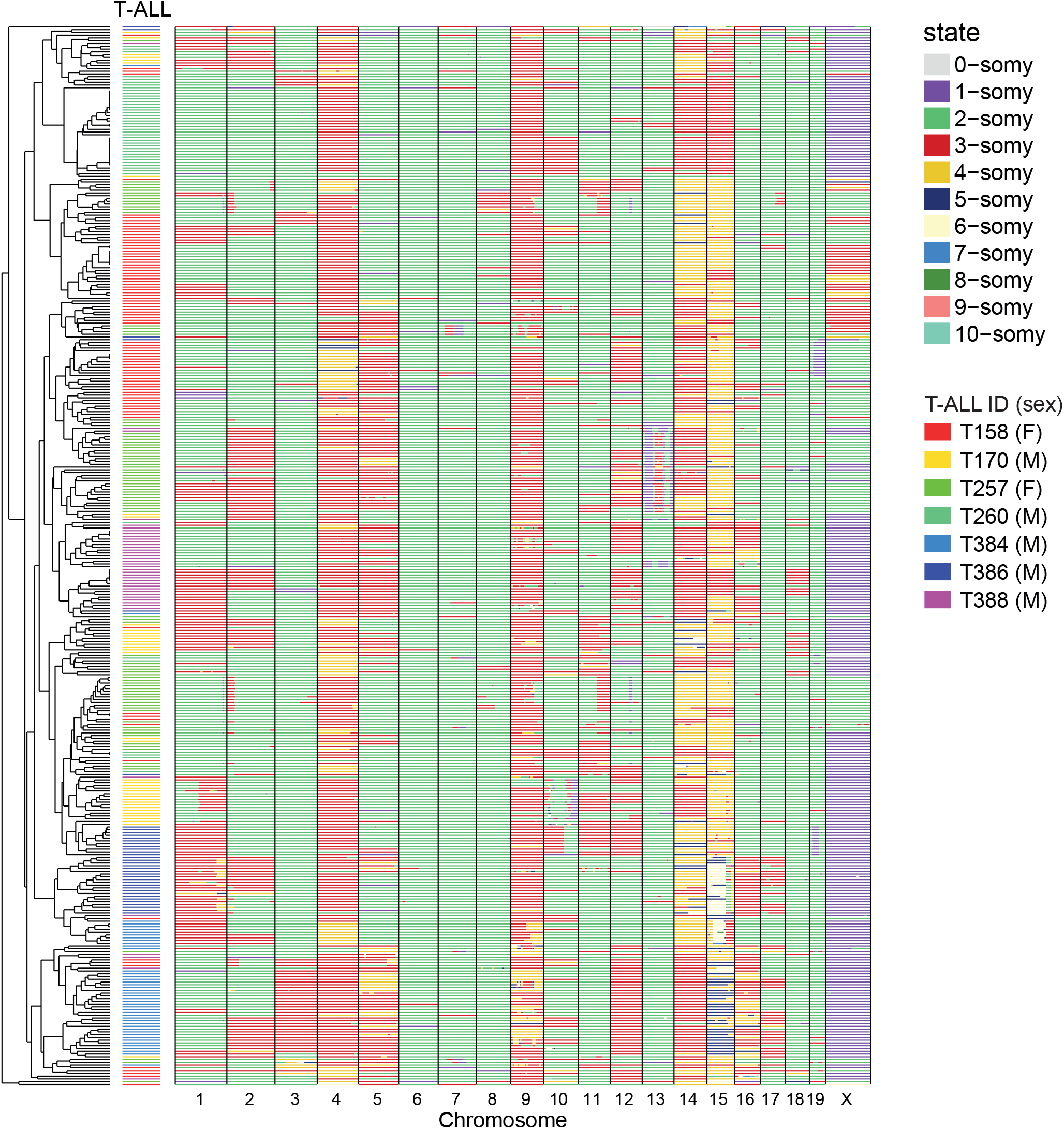
Karyotype landscape of Mps1^DK^ T-ALL. Combined scWGS data from 382 individual lymphoma cells from 7 independent T-ALLs, clustered by karyotype similarity, showing the consensus karyotype for CIN-driven T-ALL cells. Cells are clustered by karyotype similarity.

**Figure S4:**
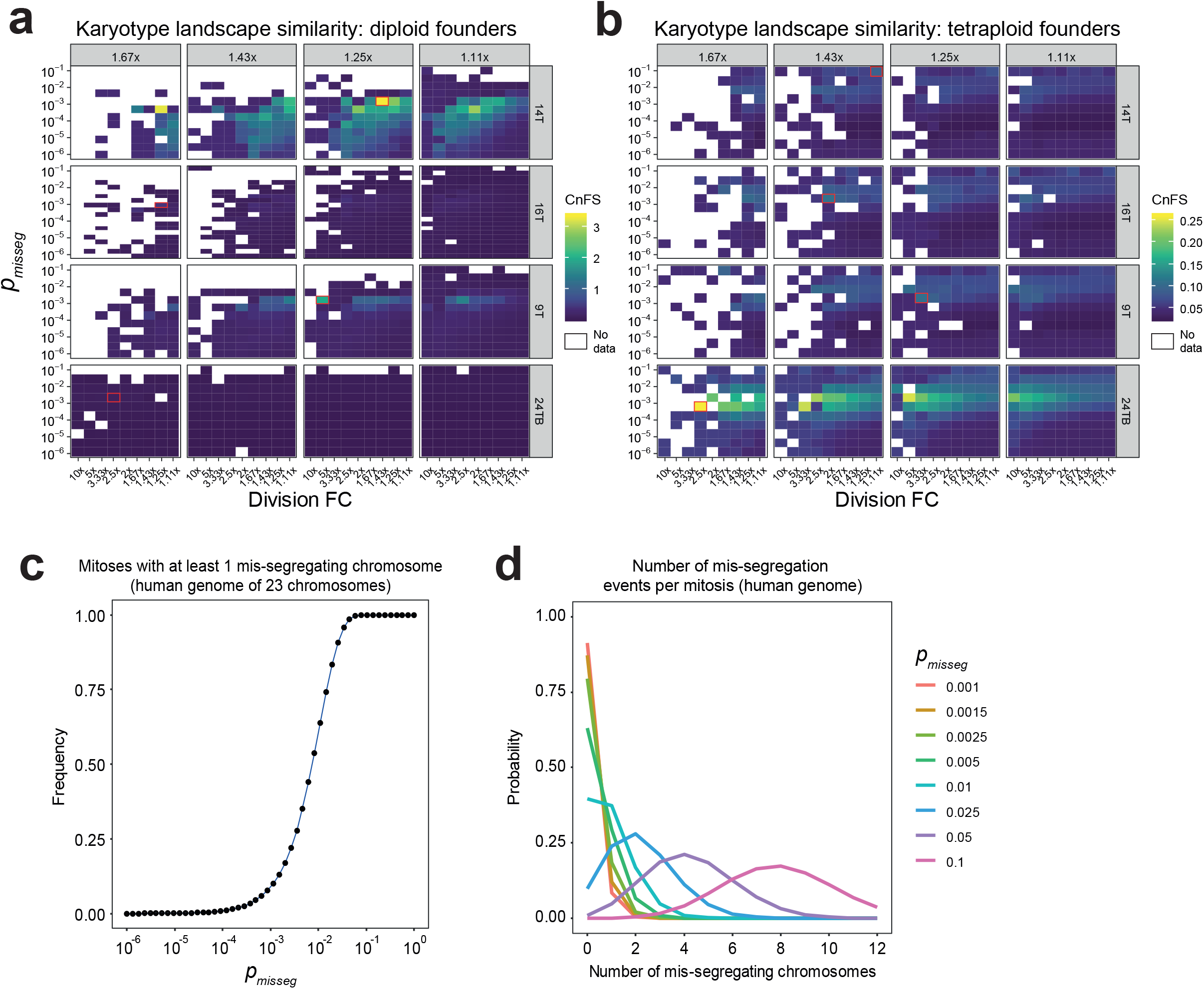
Human karyotype mis-segregation dynamics and *in vitro* mis-segregation rates. **a.** Karyotype landscape similarity quantified by CnFS for human CRC organoid simulations using diploid founders, optimal division and survival FCs and *p_misseg_* highlighted with a red box. **b.** Same as in panel a, using tetraploid founders instead. **c**. Fraction of mitoses resulting in at least 1 mis-segregating chromosome as a function of *p_misseg_* for human karyotypes. Data represents the mean±SD of 25,000 mitoses per value of *p_misseg_*. **d.** The observed frequency of chromosome mis-segregation events per mitosis for human cells. Data represent mean of 25,000 mitoses.

